# Sustained GnRH Agonism Alters Endocrine Dynamics and Pubertal Progression in Juvenile Rats

**DOI:** 10.64898/2026.06.26.734882

**Authors:** Thomas Niepsuj, Rithika Nurani, Gabriela de Faria Oliveira, Aimee K. Johnson, Amber T. Nguyen, Anna Jesch, Kristin M. Ebert, Walid Farhat, Joan S. Jorgensen, Anthony P. Auger

## Abstract

**Purpose:** Gonadotropin releasing hormone (GnRH) agonists are clinically used to delay pubertal progression by suppressing the hypothalamic-pituitary-gonadal (HPG) axis. While GnRH agonists have long been used clinically, the developmental characterization of HPG axis suppression during puberty remains incompletely understood. Thus, we examined the effects of GnRH receptor agonism in juvenile rats.

**Hypothesis:** Sustained GnRH receptor agonism will result in lower gonadal mass, blunt peripheral pubertal landmarks, and alter hormonal signaling dynamics within the HPG axis.

**Methods:** Animals received a single injection of extended-release leuprolide acetate depot (LA) or vehicle control on postnatal day (PND) 23. Animals were assessed for body mass and peripheral markers of puberty. On PND 44, animals were euthanized and tissues were evaluated to assess additional markers of pubertal maturation, pituitary gene transcript levels, and hormone concentrations in serum and gonads.

**Results:** In females, LA treatment resulted in a smaller gonad size, increased body mass, and less vaginal openings. In males, LA treatment resulted in smaller gonads but did not significantly alter body mass or preputial separation. In the pituitary, LA-treated rats had lower *Gnrhr*, *Fshb*, and *Lhb* transcript levels regardless of sex, while females exhibited higher *Cga* and *Nr5a1*. Serum FSH and ACTH were lower in LA-treated animals, and treated females also had lower progestins and androstenedione, and higher LH.

**Conclusions:** LA treatment reduced aspects of pubertal maturation and HPG axis output, with sex specific outcomes. These findings highlight the need for integrated, multi-level approaches to understand how altered GnRH signaling impacts pubertal and long-term physiology.

## Introduction

Puberty is characterized by a coordinated, multidimensional maturation of endocrine axes, organ systems, epigenetic programs, and neural circuits that together, transition the organism from a non-reproductive juvenile to a reproductively competent adult (1, 2, 3, 4). Hormones drive pubertal maturation through organizational effects that shape masculinized and feminized phenotypes that can be further activated to influence physiological and behavioral function (5, 6). At the neuroendocrine level, various pubertal triggers (7) activate endogenous pulses of Gonadotropin Releasing Hormone (GnRH) in the hypothalamus. The GnRH pulses then act in the anterior pituitary to drive rhythmic release of gonadotropins: luteinizing hormone (LH) and follicle-stimulating hormone (FSH) (8, 9). These gonadotropins can then bind ligand-specific receptors in the gonads to stimulate growth and gonadal hormone production, including estrogens and androgens. This process drives endogenous puberty and secondary sex characteristic maturation (10).

Over 4% of adolescents experience disruptions to typical puberty (11) and its timing is associated with various long-term health outcomes (12); therefore, optimizing pubertal initiation becomes important. Clinically, GnRH analogues can be used to delay pubertal timing in the cases of precocious puberty or gender dysphoria (13, 14, 15, 16). GnRH analogues modulate the hypothalamic-pituitary-gonad (HPG) axis by agonizing GnRH receptors and effectively reducing the hormonally driven maturation of secondary sex characteristics. Despite widespread use, the integrated effects of sustained GnRH receptor agonism and transient HPG axis suppression during puberty remain incompletely characterized.

Animal models serve as a critical bridge between mechanistic endocrine biology and clinical investigation in pediatric populations; allowing the field to address important questions regarding the consequences of altered pubertal timing. The juvenile rat provides an effective and experimentally controlled model with a conserved HPG axis, allowing for focused mechanistic and translational investigation. Early work determined GnRH agonist treatment could be used to delay peripheral markers of puberty (17) in rodents. Subsequent work has frequently employed daily GnRH agonists to further characterize specific markers of HPG/pubertal output. Extended-release depot formulations, however, are more common in clinical practice (14, 15, 16), and remain underutilized in animal models. This limits our understanding of how altered pubertal timing using extended-release leuprolide acetate (LA) depot shapes organismal biology. As a result, an integrated understanding of how sustained GnRH receptor agonism shapes coordinated endocrine, molecular, and physiological outcomes during puberty is lacking. Here, we address this gap by combining measures of HPG axis function with a clinically relevant LA depot formulation in prepubertal male and female rats. We hypothesize that sustained GnRH receptor agonism will: (1) reduce the size of sex steroid responsive organs, importantly the gonads; (2) delay or blunt peripheral pubertal landmarks; and (3) alter hormonal signaling dynamics within the HPG axis. Together, this framework more comprehensively characterizes the impact of GnRH agonism on endocrine systems during pubertal development in adolescent rats.

## Materials and Methods

### Animals

All experimental procedures were performed and approved by the University of Wisconsin–Madison Animal Care and Use Committee. On postnatal day (PND) 21, 48 juvenile Sprague-Dawley Rats (24 males, 24 females) were shipped from Charles River Laboratories (Massachusetts, United States) to the University of Wisconsin–Madison Department of Psychology vivarium (Madison, Wisconsin). Animals were group housed in polycarbonate cages with four animals per cage. Cages consisted of same-treatment and same-sex cage mates. Sex was determined by anatomical assessment of anogenital distance and presence of testes. Animals were kept on a 12:12 light/dark cycle, with food (Lab Diet: Laboratory Rodent Diet 5001) and water ad libitum. Cages were fitted with wood shavings (Nepco: Shredded Aspen Shavings) for bedding and nylon bones (1 per rat) for enrichment. Body mass of animals were taken on PND 23, 34 and 44.

### Drug Treatment

Animals were given 24 hours following arrival to acclimate to our facility. On PND 23, rodents were taken out of their home cage six hours into their dark cycle to receive treatment of leuprolide acetate as a depot suspension (LA) or vehicle control. All animals were weight matched to ∼55 grams (±10 grams). PND 23 was chosen as a juvenile, early peripubertal time point, prior to peripheral markers of puberty. All injections were performed subcutaneously, and injectors were unaware of the treatment condition.

The vehicle was prepared according to the manufacturer’s formulation and consisted of 5 mg/mL carboxymethylcellulose, 50 mg/mL D-mannitol, and 0.94 µL/mL polysorbate 80 in deionized water. Commercially available Lupron Depot (3.75 mg per syringe; NDC 0074-3641-03) was reconstituted according to the manufacturer’s instructions to generate a leuprolide acetate depot suspension, which was subsequently diluted with vehicle to facilitate accurate dosing. Treated animals received a single subcutaneous injection of 0.24 mL suspension (0.6 mg leuprolide acetate/rat), whereas control animals received an equivalent subcutaneous volume of vehicle.

Dosing was calculated based on clinical weight-based recommendations and projected body weight at the anticipated time of euthanasia rather than body weight at the time of administration; as juvenile Sprague–Dawley rats undergo rapid growth during the treatment interval. Projected growth trajectories were estimated using vendor-provided growth curve data (Charles River Laboratories). The final dose (0.6 mg/rat) was selected to ensure sustained exposure throughout the study period and is comparable to doses previously reported in rats when normalized to body weight (18, 19).

### Tissue Collection

On PND 44, rats were anesthetized with isoflurane and immediately decapitated. Trunk blood was collected in Vacuette® 4 mL CAT serum separator tubes (Greiner Bio-One, 456292P) and centrifuged at 3,000–5,000 RPM for 10 min at 25 °C using an IEC Clinical Centrifuge (model CL, serial 72938M, 115 VAC, 50/60 Hz, IEC). Serum was aliquoted and stored at −20 °C until hormone analysis. Pituitaries were removed, flash-frozen in 2-methylbutane on dry ice, and stored at −80 °C until processed. Gonads and seminal vesicles were removed and weighed; gonads were similarly flash-frozen and stored at −80 °C. Penises were dissected, placed in 4% PFA overnight and then processed for H&E staining by the University of Wisconsin-Madison Experimental Animal Pathology Laboratory (EAPL).

### Peripheral Markers of Pubertal Development

Vaginal opening (20) and balanopreputial, or preputial, separation (21) were assessed as peripheral markers of puberty to monitor drug effectiveness. Vaginal opening was checked on PNDs 23, 34, and 44. Preputial separation is typically completed by PND 44 (22), and was assessed histologically at euthanasia. To standardize preputial separation analysis, the presence of the baculum and hyaline cartilage was confirmed in each section to verify a medial parasagittal plane of the penis, using histological anatomy resources (23, 24). Preputial separation was assessed by measuring the length of cornified papillae along the preputial lining relative to the basal lining of the prepuce (25). Examiners were unaware of the treatment group for all analyses.

### Pituitary Quantitative Polymerase Chain Reaction

Pituitary samples were homogenized via sonication, and total RNA was collected using a PureLink RNA Mini Kit (Thermo Fisher Scientific; catalog No. 12183018A) according to manufacturer directions. RNA purity and concentrations were measured using a NanoDrop 2000 spectrophotometer (Thermo Fisher Scientific; catalog No. ND-2000LAPTOP) and diluted such that all samples had a standard concentration of 20ng/uL. 100-200ng of RNA was converted to complementary DNA (cDNA) using the High-Capacity RNA-to-cDNA Kit (Thermo Fisher Scientific; catalog No. 4387406) using manufacturer directions. Real-time quantitative polymerase chain reaction (qPCR) was conducted using the CFX Opus 96 real-time PCR system, and cDNA was amplified with PowerUp SYBR Green Master Mix (Thermo Fisher Scientific; catalog No. A25741). After amplification, only reactions with 100% efficiency (±10%) and acceptable dissociation curves confirming product purity were included in the analysis.

All genes of interest were normalized to a housekeeping gene, ribosomal protein L13A (*Rpl13a*; NM_173340): forward primer AGCAGCTCTTGAGGCTAAGG and reverse primer GGGTTCACACCAAGAGTCCA using the 2^−ΔΔ^Ct method as published previously (26). Primers for genes of interest were purchased from IDT DNA technologies as predesigned PrimeTime™ PCR Primers. Primer details, as given by the IDT specification sheet, are summarized in table 1.

**Table 1.**
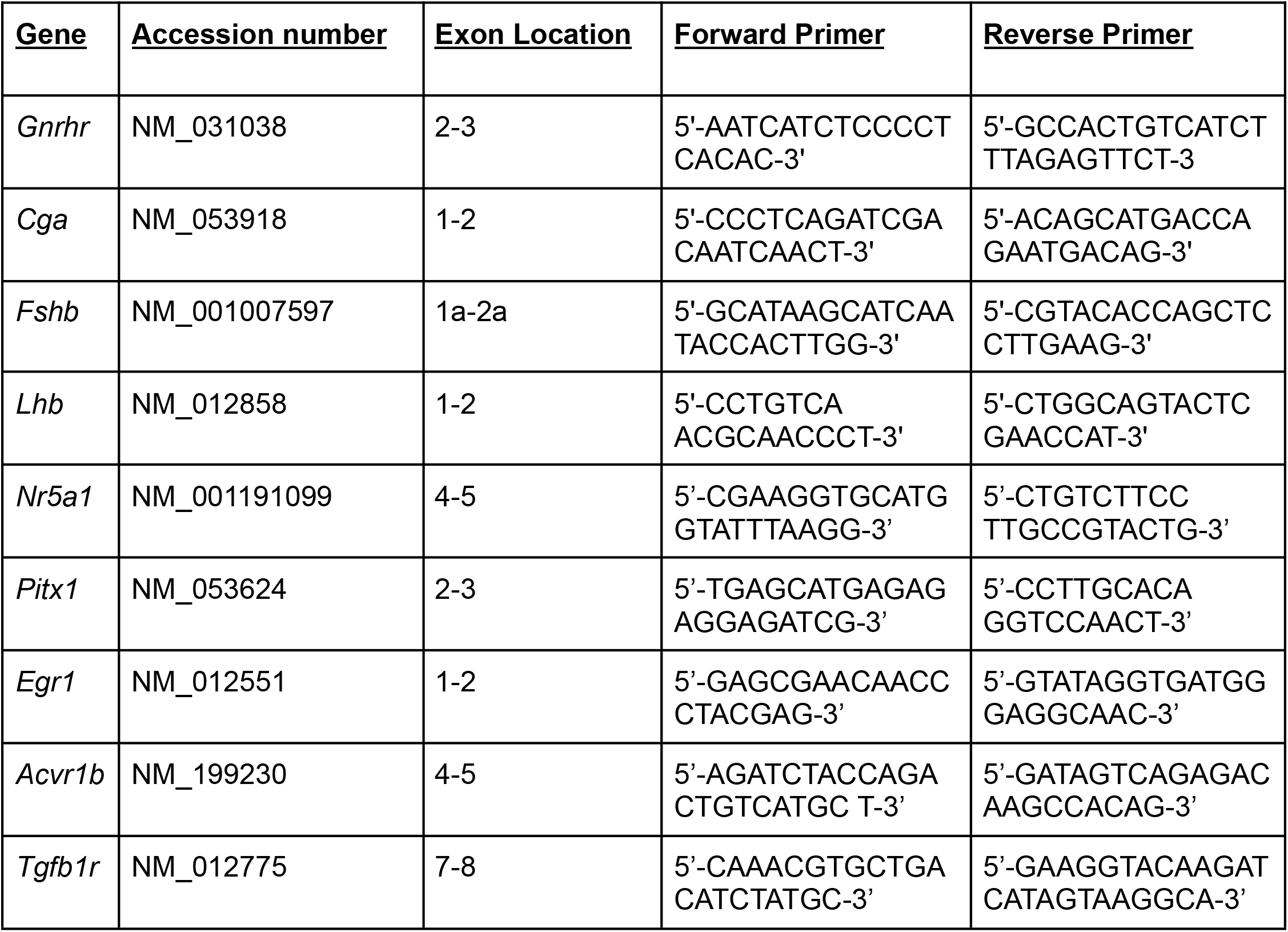
Pituitary Transcripts. All analyzed via qPCR. The table includes accession numbers, exon locations, and primer sequences.

### Hormones

Serum and gonad samples were submitted to the Wisconsin National Primate Research Center (WNPRC) Assay Services for hormone analysis. Serum steroid hormones: corticosterone, progesterone, 17-hydroxyprogesterone, estradiol, estrone, androstenedione and testosterone were measured by liquid chromatography–tandem mass spectrometry (LC-MS/MS) using a Sciex 6500+ Mass Spectrometer. Intraovarian estradiol and progesterone and intratesticular testosterone were similarly measured from homogenates of whole ovaries and representative testicular samples (∼1/8–1/16 of total tissue mass). All analytes were measured in duplicate using 200µL serum (n=10/group) or 50mg tissue (n=6/group) per sample.

Adrenocorticotropin (ACTH), brain-derived neurotrophic factor (BDNF), follicle-stimulating hormone (FSH), growth hormone (GH), luteinizing hormone (LH), prolactin, and thyrotropin stimulating hormone (TSH) were measured using a MILLIPLEX MAP Rat Pituitary Magnetic Bead Panel—Endocrine Multiplex Assay (catalog No. RPTMAG-86 K, RRID:AB_2716840). The multiplex panel was performed according to the manufacturer’s instructions using a Bio-Plex 200 system. Serum samples were diluted 1:3 by combining 25 µL serum with 50 µL assay diluent (serum matrix). Subsequently, 25 µL of the diluted sample was loaded into each well and analyzed in duplicate (n = 10/group). Additionally, myostatin concentrations were measured in duplicate using the Quantikine GDF-8/Myostatin Immunoassay (Catalog No. DGDF80, RRID: AB_2905547), according to the manufacturer’s instructions; on an Accuris Instruments SmartReader 96 Absorbance Plate Reader (100uL/well, n = 10/group). Samples that were below the level of detectability were set to the lowest detectable level to allow for statistical analysis of hormones; notably FSH, progesterone, 17-hydroxyprogesterone, and female testosterone.

### Data Analysis and Manuscript Preparation

All data were analyzed using GraphPad Prism 10 Software version 10.5.0 (673) for MacOS (GraphPad Software, San Diego, California). Outliers were identified using GraphPad Prism’s Grubbs’ test (α = 0.05), with a maximum of one outlier removed per group for each dataset. Samples with technical assay failures were excluded from analysis. Data with two group comparisons were analyzed using Welch’s t-test. Fisher’s exact test was additionally run on binary data sets, such as vaginal opening. Four group comparisons were analyzed using 2-way analysis of variance (ANOVA) with main effects examining treatment and sex. When interactions between treatment x sex were detected, comparisons were followed by Tukey’ post-hoc analysis. All figures express mean ± SEM and all statistical significance was set at a p-value less than or equal to 0.05. The authors used ChatGPT (OpenAI, GPT-5) for language editing and improving sentence structure, formatting and grammar. All data, analyses, and conclusions are the original work of the authors.

## Results

### Body and Reproductive Organ Mass

Animals were tracked longitudinally for body mass on PND 23, 34 and 44 (Fig. 1A). All animals were weight matched on PND 23 to ∼55g. Sex differences in body mass were present by PND34 (2-way ANOVA main effect of sex, p = 0.0167; Data not shown). At PND 44, both treatment and sex differences were observed (interaction of sex x genotype, p = 0.0020; Fig. 1B). Treated females had higher body mass than control females (Fig. 1A; Tukey post hoc, p = 0.0365) that eliminated sex differences between treated females and males. At euthanasia, treated ovaries, testis and seminal vesicles had lower masses than controls (data not shown), with lower masses maintained when normalized to body mass (Fig. 1C-E).

**Figure 1.**
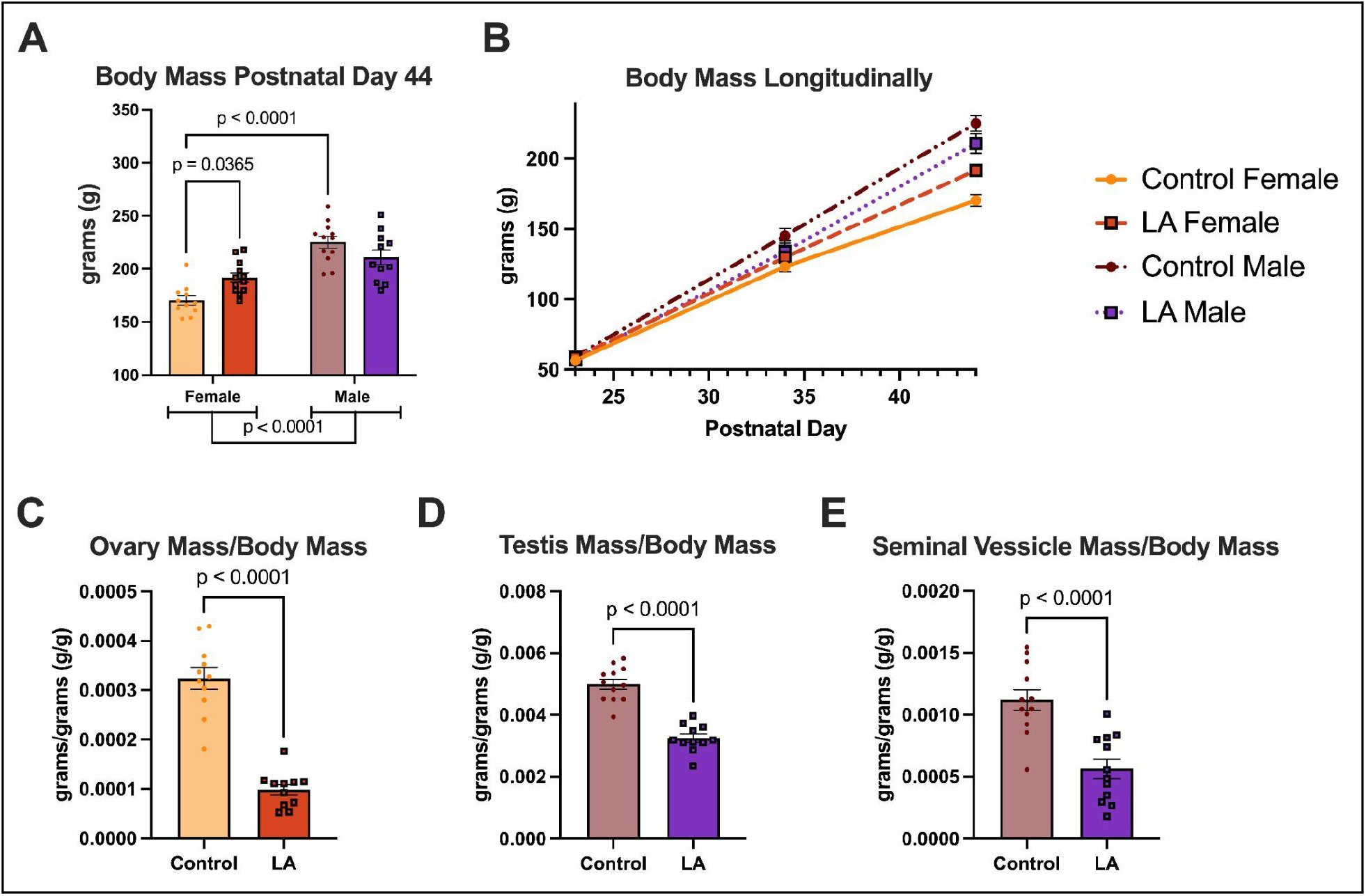
Body and reproductive organ mass. **A)** Body mass of rodents at euthanasia, postnatal day 44. **B)** Longitudinal body mass rats treated with vehicle control or .6mg/rat of extended release leuprolide acetate (LA). All groups were weight matched on treatment date, PND 23. Gonad and accessory organ mass, when normalized to body mass {**C)** Ovary, **D)** Testis, **E)** Seminal vesicle. All significant p values (p < 0.05) displayed within graphs. Data represented as mean ± SEM and all figures n=12/group. [Top Right] Figure Legend: Yellow circles & solid line, control females; orange squares & dashed line, treated females; maroon circles & dash-dot line, control males; purple squares & dotted line, treated males.

### Peripheral Markers of Puberty

All females, regardless of treatment, had no vaginal opening at PND 23 (n=12/group, data not shown). At PND 34, 75% of LA depot treated females did not have vaginal opening, which persisted through PND 44 (Fig. 2A, Fisher’s exact test, p = 0.0003). Preputial separation was assessed under histological examination (n=6/group) at PND 44. LA depot treated males were comparable to controls, but more likely to have incomplete separation (Fig. 2B, p=0.1422).

**Figure 2.**
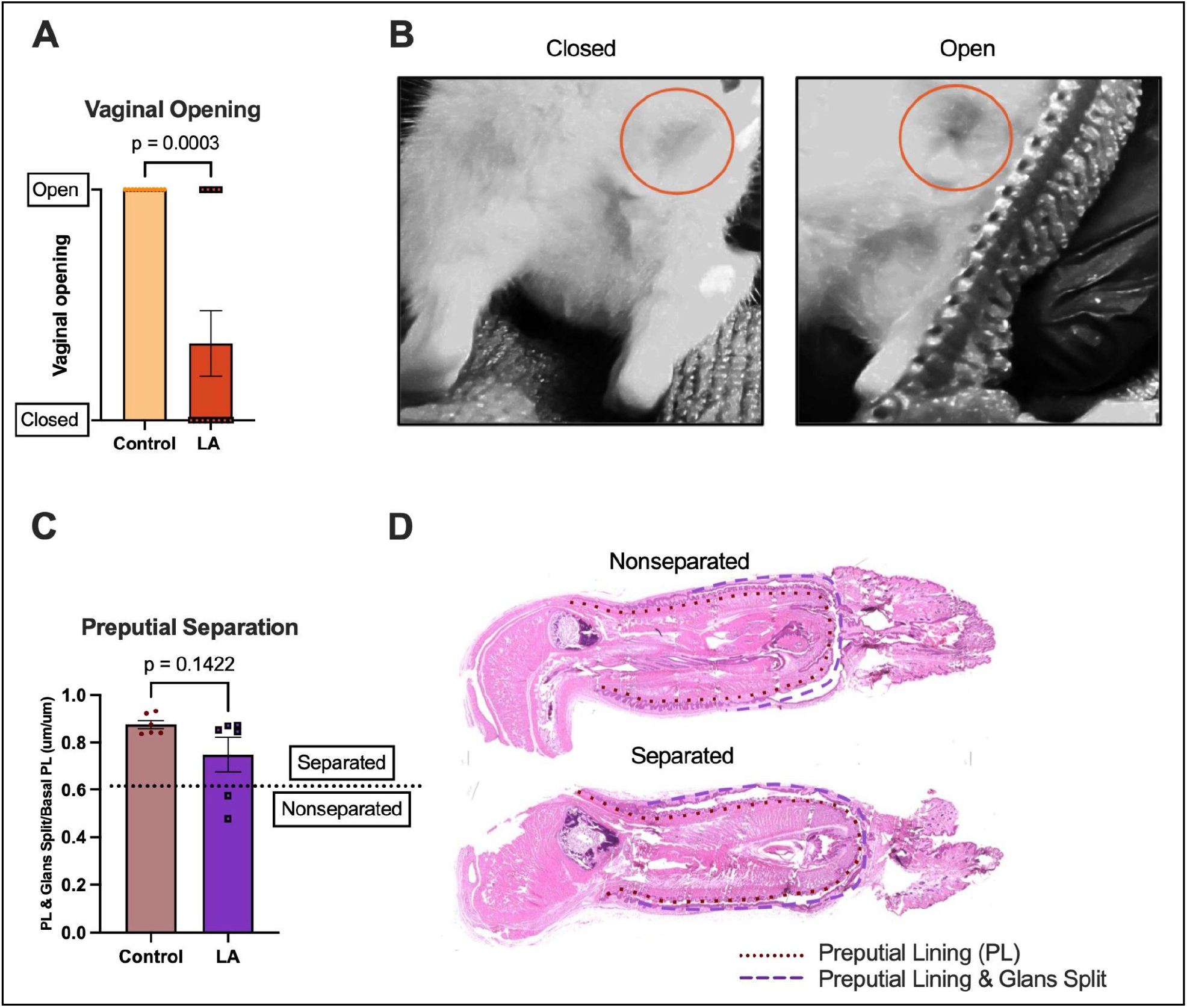
Peripheral markers of rat puberty. **A)** Female peripheral marker of puberty, vaginal opening, checked on both PND 34 and 44 (n=12/group). **B)** Representative photos of rats with closed [left] and opened [right] vagina. **C)** Comparison of histological preputial separation on PND 44. Data represents the length of the glans split from the basal preputial lining (PL) over the length of the basal PL (n=6/group). The dashed line indicates an operational threshold used to categorize animals as having completed or not completed preputial separation. **D)** Representative image of H&E stained rat penis, including a nonseparated [top] and separated [bottom] phenotype. P values are displayed within graphs, with data represented as mean ± SEM. Yellow circles, control females; orange squares, treated females; maroon circles, control males; purple squares, treated males.

**Figure 3.**
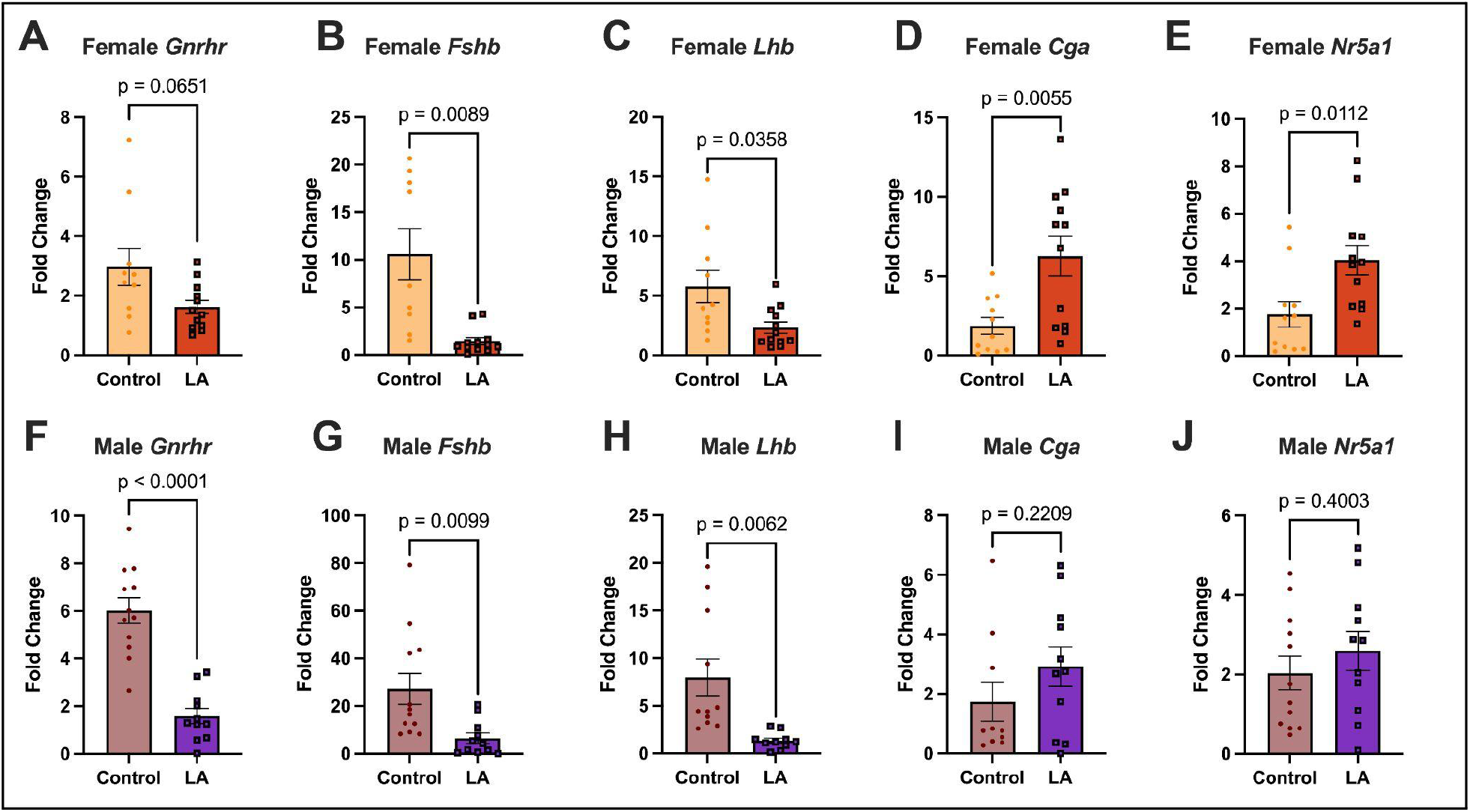
Pituitary gene transcripts of gonatrope signaling. [Top row, A-E] Female pituitary gene expression. **A)** *Gnrhr* **B)** *Fshb* **C)** *Lhb* **D)** *Cga* **E)** *Nr5a1*. [Bottom row, F-J] Male pituitary gene expression. **F)** *Gnrhr* **G)** *Fshb* **H)** *Lhb* **I)** *Cga* **J)** *Nr5a1*. P values are displayed within graphs with data represented as mean ± SEM; all graphs n=12/group. Yellow circles, control females; orange squares, treated females; maroon circles, control males; purple squares, treated males.

### Pituitary Transcripts

Pituitary transcripts involved in gonadotrope signaling were assessed on PND 44. Females and males treated with LA depot exhibited differential transcript outputs. Female transcript levels of *Gnrhr* were not significantly different (Fig. 2A, p = 0.0651), while male *Gnrhr* transcript levels were significantly lower than controls (Fig. 2F, p < 0.0001). Both treated female and male pituitaries harbored significantly less transcripts for gonadotropin specifying subunits *Fshb* and *Lhb* (Fig. 2B, p = 0.0089; Fig. 2C, p = 0.0358; Fig. 2G, p = 0.0099; Fig. 2H, p = 0.0062). The transcript levels of the common alpha subunit, *Cga,* and *Nr5a1* were significantly higher in treated females (Fig. 2D, p = 0055; Fig. 2E, p = 0.0112) with no change observed in treated males (Fig. 2I, p = 0.2209; Fig. 2J, p = 0.4003). No differences were observed in *Acvr1b*, *Egr1*, *Ptx1*, or *Tgfb1r* pituitary transcripts (data not shown).

### Hormones

Serum was taken to assess peripheral hormones. LA treated animals had a lower concentration of FSH in females (Fig. 4A, p = 0.0179) and males (Fig. 4F, p = 0.0005). Similarly, serum progestins and androstenedione concentrations were lower in treated females (Fig. 4C, p = 0.0058; Fig. 4D, p = 0.0224; Fig. 4E, p = 0.0027). Serum LH, however, was higher in treated females (Fig. 4B, p = 0.0079). No other significant treatment-versus-control comparisons within either sex were observed by Welch’s t-test. To evaluate the effects of treatment and sex across all animals, serum hormone concentrations were also analyzed using two-way ANOVA. Treated animals had higher ACTH than controls (Fig. 5F, p = 0.0299). Sex differences were observed in corticosterone (Fig. 5A, p = 0.0128), prolactin (Fig. 5B, p = 0.0184), androstenedione (Fig. 5E, p = 0.0004), testosterone (Fig. 5D, p < 0.001) and BDNF (Fig. 5H, p = 0.0018). No differences were observed in growth hormone, TSH, or myostatin (data not shown). Estradiol and estrone were below the level of detectability in males and females, so no meaningful analysis was conducted. No significant differences were observed in intragonadal steroids (Fig. 6).

**Figure 4.**
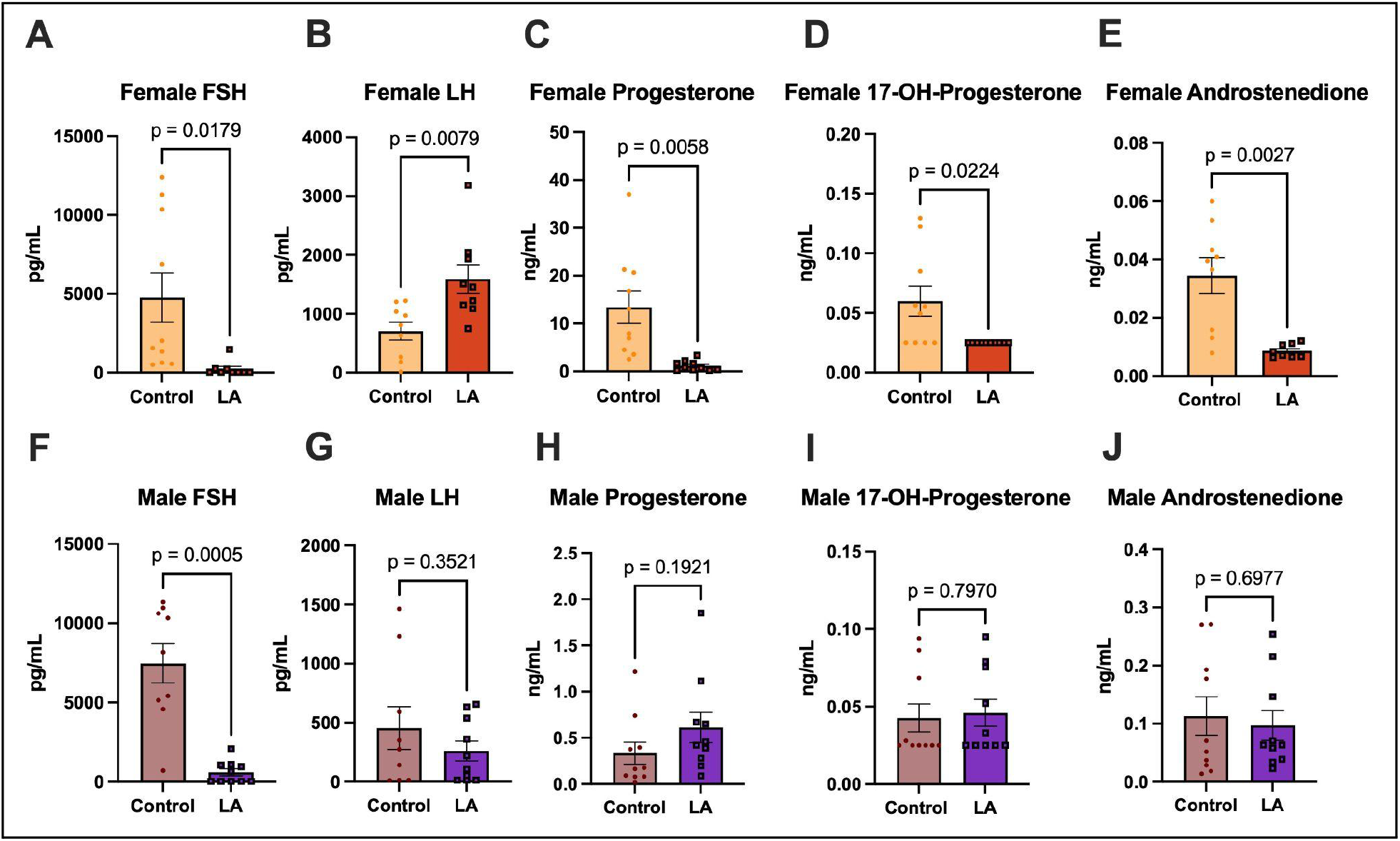
Serum Hormones. [Top row, A-E] Female serum hormones. [Bottom row, F-J] Male serum hormones. **A&F)** Follicle-Stimulating Hormone (FSH) **B&G)** Luteinizing Hormone (LH) **C&H)** Progesterone **D&I)** 17-hydroxyprogesterone (17-OH Progesterone) **E&J)** Androstenedione. P values are displayed within graphs with data represented as mean ± SEM; all graphs n=10/group, unless a single grubbs outlier is removed.

**Figure 5.**
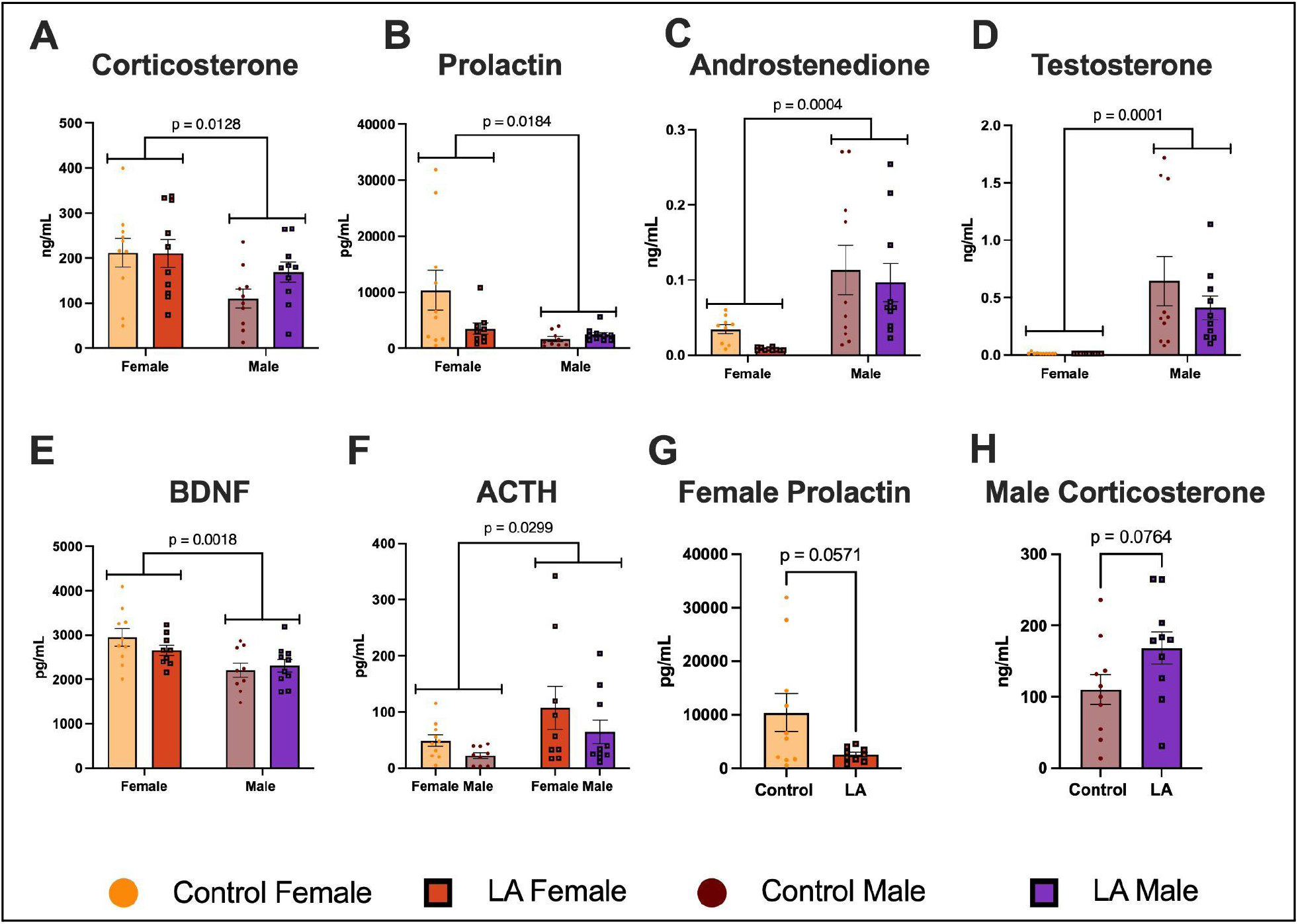
Additional Serum Hormone Comparisons. **A)** Corticosterone **B)** Prolactin **C)** Androstenedione **D)** Testosterone **E)** Brain Derived Neurotropic Factor (BDNF) **F)** Adrenocorticotropin (ACTH) **G)** Female Prolactin **H)** Male Corticosterone. P values are displayed within graphs with data represented as mean ± SEM; all graphs n=10/group, unless a single grubbs outlier is removed. [Bottom] Figure Legend: Yellow circles, control females; orange squares, treated females; maroon circles, control males; purple squares, treated males.

**Figure 6.**
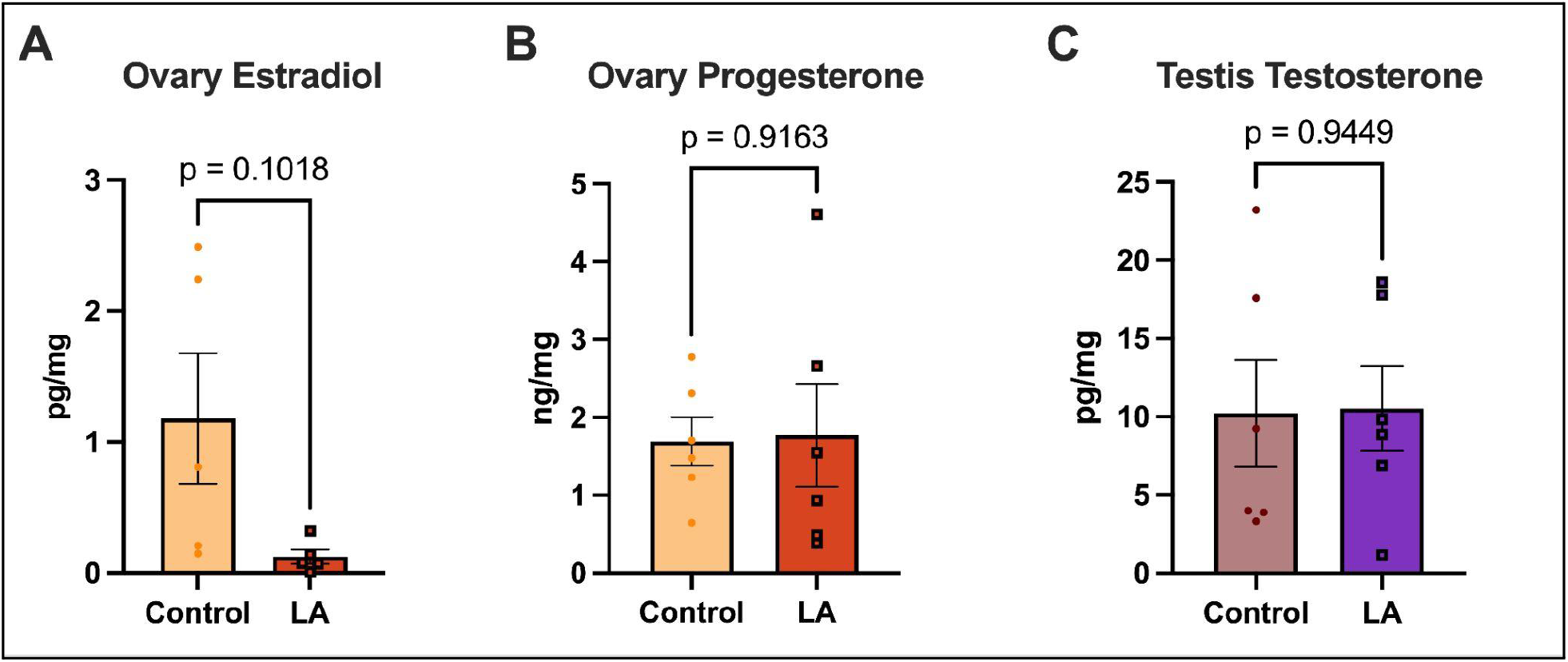
Intragonadal Steroid Measurements. P values are displayed within graphs, with data represented as mean ± SEM, all graphs n=6/group, unless a single grubbs outlier is removed. Yellow circles, control females; orange squares, treated females; maroon circles, control males; purple squares, treated males.

## Discussion

This study integrated multiple levels of analyses, including body and tissue mass, peripheral markers of puberty, gene expression, and hormone concentrations to evaluate the influence of GnRH agonism in juvenile rats. We found that LA depot treatment early in adolescent development produced changes in mass, pituitary gene expression, and hormone concentrations, with sex specific outcomes. Collectively, these results indicate that adolescent GnRH agonism influences multiple coordinated levels of endocrine regulation beyond peripheral markers of puberty.

### Body and Reproductive Organ Mass

Growth is a hallmark of pubertal maturation and is regulated by circulating steroids and metabolic cues (27, 28). Sex differences in body mass and composition emerge during puberty and are largely driven by gonadal steroid signaling (29, 30). Consistent with the well-established role of estrogen in modulating female body composition and limiting weight gain (30, 31, 32), our control females weighed less than control males. In contrast, LA depot treatment eliminated these sex differences, with treated females having a higher body mass that approached control males, and no significant differences observed between treated females and treated males. This pattern of eliminated sex differences is consistent with gonadectomized animals (33, 34). While we did not directly assess adiposity or fat distribution in these animals, several rodent studies suggest that suppression of ovarian steroids can influence feeding behavior and metabolism, promoting weight gain (30, 32). While we did not observe differences in estrogen output in LA treated animals at PND 44, it is possible that we missed the timing of estrogen suppression that allowed for the male-typical increase in body weight. Notably, the higher body mass, which is associated with accelerated pubertal progression, did not influence vaginal opening in LA depot treated females. These findings highlight a dissociation between somatic growth and peripheral pubertal progression under GnRH agonism, as higher body mass alone was insufficient to advance markers of sexual maturation. This suggests that somatic growth, steroid hormone signaling, and reproductive axis activation are coordinated but can be differentially regulated under pharmacologic perturbation.

Clinically, adolescents treated with GnRH agonists often exhibit higher body mass index and adiposity (35, 36, 37). This is consistent with altered metabolic regulation during suppression of gonadal steroid production in the juvenile period, particularly in females. Together, these findings parallel clinical observations in which GnRH agonist treatment is associated with increased body mass and adiposity, particularly in females, suggesting conserved effects of hormonal modulation on metabolic regulation across species. Human reports, however, are not all conclusive (38).

Similarly, reproductive organ maturation serves as an additional marker of pubertal development. As an index of this process, reproductive organ mass was assessed and markedly suppressed in LA depot treated animals, regardless of sex. Gonadal size is known to remain smaller relative to controls under pharmacologic suppression of the HPG axis via various GnRH agonists, (39, 40, 41, 42, 43, 44, 45, 46, 47). More specifically, daily agonism by leuprolide acetate blocked the seminal vesicle mass increases, but had a lesser effect on testis mass (48, 49, 50). Notably, daily injections of leuprolide acetate during adolescence had no influence on ovarian (51, 52) or testis mass (53) following a washout period.

### Peripheral Markers of Puberty

Vaginal opening and preputial separation are often considered the hallmark indicators of peripheral puberty in rats. We used these metrics to assess an aspect of pubertal maturation and found sex specific results.

Vaginal opening is closely related to changes in sex hormones and the ovulatory process (54). This process is strain dependent and typically occurs in Sprague-Dawley rats around PND 32 (55). We assessed vaginal opening shortly after this time point and at PND 44, the euthanasia time point. The majority of animals treated with LA depot on PND 23 had no vaginal opening by PND 44. Although vaginal opening is widely used as a marker of sexual maturation in females and is accelerated by activation of the HPG axis (56), classic early development ovariectomy studies demonstrated that it can occur independently of peripubertal ovarian output (57); suggesting that additional extra-ovarian sources contribute to vaginal opening. GnRH agonism has been shown to both accelerate and delay vaginal opening, depending on the treatment paradigm (58). In our model, LA depot prevented vaginal opening in the majority of animals, indicating that this upstream modulation of HPG output is more effective at blocking vaginal opening than early ovariectomy alone. Importantly, vaginal opening is not the sole indicator of sexual maturity in females, with first estrus and first ovulation being among additional maturity indicators (59). Future research should assess these markers longitudinally following the administration of LA depot.

Preputial separation is a male peripheral marker of puberty, which is characterized by the prepuce separating from the glans penis (60); a process driven by low levels of androgens (25) and typically completed by PND 44 (55). While vaginal opening can be assessed visually, preputial separation is more difficult to evaluate, as distinguishing complete from incomplete separation is challenging and repeated examinations may accelerate the process (61). To more quantitatively assess preputial separation, histological analysis of the cannulation of the preputial space was measured by comparing the length of the basal lining to the distance of canalized space. Although treated animals were more likely to have incomplete preputial separation, there were no significant differences. This may suggest that there was a delay in peripheral puberty in males. By study completion at PND 44, however, circulating testosterone was comparable between treated and control males, thus, the timing and/or length of treatment were insufficient to fully block the androgen threshold required for this peripheral marker of puberty.

### Gonadotrope Signaling and Gonadotropin Transcripts

The pituitary is a master mediator of endocrine action that mediates GnRH signals and peripheral hormone production. Gonadotropes are a major population of hormone-secreting cells in the anterior pituitary that produce the gonadotropins FSH and LH. These hormones are heterodimers composed of a common α-Alpha Glycoprotein Subunit (αGSU), encoded by *Cga*, and hormone-specific β-subunits encoded by *Fshb* and *Lhb*, respectively. Importantly, GnRH pulse frequency differentially induces gonadotropin production (62, 63). In the context of GnRH agonism, it is known that GnRH receptors follow a homologous desensitization from persistent GnRH signaling (64). Consistent with this, a reduction in *Gnrhr* mRNA is observed following GnRH agonism via depot administration in juvenile rats (19), which is recapitulated in our data set, with more pronounced effects in males. In contrast, daily agonism in juvenile rats has been shown to increase *Gnrhr* mRNA in the rat pituitary regardless of sex (65), highlighting a potential treatment administration or timing effect on *Gnrhr* mRNA. Further, it has been previously reported that agonism leads to a down regulation of Lhb and Fshb subunit transcript expression (18, 66, 67), which is supported by our results in both males and females. Kim, Swerdloff & Bhasin, however, observed an increase in *Lhb* mRNA after four weeks of daily leuprolide acetate treatment (68); highlighting that timing and dosing schedule of GnRH agonism differently modulates HPG axis output.

Although *Gnrhr*, *Lhb*, and *Fshb* mRNA levels follow expected patterns, we saw no influence in *Cga* expression in males and an increase in females. It is known that stimulation of the GnRH receptor can upregulate *Cga* expression *in vitro* (69), and GnRH agonism has upregulated *Cga in vivo* (66, 68). These observations support our data. Additionally, it has been similarly reported that human females have an increase in free alpha subunit protein levels towards the end of an extended LA treatment, with beta subunit protein levels being undetectable (70). Similarly, *Nr5a1* (encoding steroidogenic factor 1, a driver of gonadotrope cell differentiation) was upregulated in the same female-specific pattern. *Nr5a1* is a critical regulator of reproductive maturation in rodents (71, 72); thus, its elevated expression in females may reflect ongoing pituitary gonadotrope differentiation and activation of pathways associated with sexual maturation. This female specific increase in gonadotrope differentiation may also explain increased output of *Cga*. Notably, these data are derived from a single time point and therefore do not capture the pulsatile dynamics of GnRH activity and gonadotrope transcription. They do, however, complement existing datasets and support drug efficacy at the pituitary transcript level, as evidenced by reduced expression of genes involved in gonadotropin production.

### Hormones

Pituitary and steroid output are additional marks of HPG maturation and LA effectiveness. Continuous GnRH agonism has been reported to downregulate Gnrhr transcripts in vitro; thereby decreasing gonadotropin subunit expression and hormone output (73). In alignment with the down regulation of pituitary *Fshb* transcript levels and FSH’s constitutive release (74), we measured less serum FSH, which has been similarly reported with daily leuprolide acetate (26). LH output, however, is more tightly regulated by GnRH (74), and had a more nuanced phenotype. As predicted, LA depot administration resulted in less pituitary Lhb transcript levels in both sexes; however, serum LH was unchanged in males and elevated in females. This has been similarly observed, where 14+ days of GnRH agonism reduced *Lhb* mRNA levels in rats but increased or had no effect on serum LH (18, 66). Notably the pattern of serum LH more similarly follows pituitary *Cga* transcript levels, with a significant positive correlation across all subjects (F=4.771; DFn,DFd=1,29; p=0.0372). Our findings highlight that GnRH agonism produced sex-specific effects on gonadotropin regulation during the juvenile peripubertal period.

Gonadtropins are known to stimulate the gonads to produce sex steroids. In females, serum progesterone and its downstream metabolite, 17-hydroxyprogesterone, were lower than control animals; however, intraovarian progesterone levels were comparable between groups, highlighting a reduction of progesterone output. Serum estrogens were undetectable in all groups, and intraovarian estradiol was not different between groups. In males, no differences were observed in serum LH, circulating androgens, or intratesticular testosterone levels. It was surprising to see no reduction in testosterone, as GnRH agonism is expected to suppress androgens, as previously reported under daily GnRH agonism (49, 50). Differences in dosing regimen, age of treatment initiation, or age of euthanasia may explain the lack of reported androgen suppression. Given that testosterone production is primarily stimulated by LH, the comparable androgen output suggests preserved Leydig cell function. Despite unchanged circulating and intratesticular androgen levels, testis mass was significantly reduced alongside marked suppression of FSH. Given the role of FSH in supporting testicular growth, Sertoli cell function and spermatogenesis (27, 75), the reduction in testis mass is more likely attributable to decreased spermatogenic output than impaired androgen production.

A strength of our dataset is the replication of previously reported sex differences in circulating androgens, progesterone, and corticosterone (76, 77). While several endocrine outcomes varied between our previously reported daily leuprolide acetate model (26) and current LA depot condition (including GH, TSH, and testosterone), both paradigms produced changes in progesterone, suggesting a conserved effect across treatment regimens. Notably, treated animals had increased ACTH levels, which could be a sign of heightened adrenal output. Similarly, we previously reported daily Leuprolide acetate increase males to female concentrations of corticosterone (26). Although these studies were conducted under different dosing paradigms, treatment durations, hormone collection techniques, and drug delivery methods, these findings collectively suggest that administration strategy (timing, dose, and/or pharmacokinetics) can differentially shape endocrine system responsiveness.

It is important to note that tissues and serum were collected at a single time point near the reported end of the extended-release LA depot’s efficacy. Accordingly, the present findings represent a snapshot of HPG axis activity and do not capture the temporal or pulsatile dynamics that characterize many components of this endocrine system. Consequently, transient or time-dependent changes in hormone concentrations and gene expression occurring earlier or later following treatment are not detected. Despite this, LA depot continued to delay VO and reduce serum FSH at the time of collection, indicating drug effectiveness at this stage of treatment. Ongoing studies examining additional collection time points will further characterize the temporal endocrine and molecular responses to LA depot.

### Translational Relevance

Clinically, GnRH agonists are widely used for HPG suppression, yet their broader effects beyond suppression of peripheral pubertal progression remain incompletely understood. Direct study of altered pubertal timing in children is limited by ethical and practical constraints, as well as heterogeneity in timing and patient populations. As a result, the systemic consequences of disrupted HPG axis signaling across development remain poorly defined. This gap is particularly important because pubertal hormones influence multiple systems, including somatic growth, metabolism, neural circuit maturation, and neurobehavioral development. Importantly, clinical assessment of puberty relies primarily on Tanner staging, which captures external sexual maturation but does not reflect broader endocrine or systemic changes. Thus, current clinical approaches that use only peripheral assessment of puberty may not fully capture the complexity of pubertal progression and those that combine peripheral markers along with endocrine measurements gain a more holistic insight into physiological development during adolescence.

Here, we present a controlled model of sustained GnRH agonism that enables systematic investigation of how altered pubertal timing influences endocrine dynamics and pubertal progression. This is particularly relevant to pediatric conditions involving altered pubertal timing or HPG axis disruption, including variations in sex characteristics (VSCs/DSDs), chronic childhood illness, and survivorship following gonadotoxic therapies. These findings position controlled HPG axis modulation as a translational tool for understanding how differences in pubertal timing shape long-term developmental trajectories and health outcomes across pediatric populations, beyond central precocious puberty or pubertal suppression in transgender and gender diverse youth. Ultimately, this approach may help clarify how variation in pubertal onset and hormonal exposure contributes to long-term health outcomes and inform future clinical strategies for monitoring and managing pediatric populations with altered puberty.

## Conclusions

LA depot treatment altered pubertal development in juvenile rats, demonstrated by lower gonadal mass, blunted peripheral pubertal landmarks, and modified HPG axis outputs. LA depot treatment did not, however, suppress all endocrine related systems, as male physiology, including LH and testosterone, was less influenced in the present experimental conditions. This emphasizes that there are sex differences in how GnRH receptor agonism influences juvenile rodents. From a translational standpoint, modeling LA depot in animal models provides a platform for addressing clinically relevant questions that are difficult to study in pediatric populations. Together, these data provide a framework for understanding sex-specific, multi-level responses to adolescent GnRH agonism across the HPG axis and highlight the importance of integrating endocrine and tissue-level outcomes when evaluating LA depot treatment. Future studies should address how these early-life alterations in HPG axis function influence later endocrine outcomes, including potential long-term consequences of altered pubertal timing and hormonal signaling.

## Data Availability

Some or all datasets generated during and/or analyzed during the current study are not publicly available but are available from the corresponding author on reasonable request.

